# Six percent loss of genetic variation in wild populations since the industrial revolution

**DOI:** 10.1101/488650

**Authors:** D.M. Leigh, A.P. Hendry, E. Vázquez-Domínguez, V. L. Friesen

## Abstract

Genetic variation underpins population fitness and adaptive potential^1,2^. Thus it plays a key role in any species’ probability of long-term persistence, particularly under global climate change. Genetic variation can be lost in a single generation but its replenishment may take hundreds of generations^3^. For that reason safeguarding genetic variation is considered fundamental to maintaining biodiversity, and is an Aichi Target for 2020^4^. As human activities are driving declines in many wild populations^5^, genetic variation is also likely declining. However the magnitude of ongoing genetic variation loss has not been assessed, despite its importance. Here we show a 6% decline in within-population genetic variation of wild organisms since the industrial revolution. The erosion of genetic variation has been most severe for island species, with an 18% average decline. We also identified several key taxonomic and geographic information gaps that must be urgently addressed. Our results are consistent with single time-point meta-analyses that indicated genetic variation is likely declining ^6,7^. However, our results represent the first confirmation of a global decline, and estimate of the magnitude of the genetic variation lost from wild populations.

---

Mitochondrial sequence diversity is substantially lower in geographic regions heavily affected by human activity^6^. Similarly, nuclear variation is lower in fragmented populations relative to undisturbed populations^7^. These single time-point analyses suggest that genetic variation may have been lost from many populations and is likely in global decline. However, single time-point analysis cannot disentangle contemporary declines in variation from historically low values caused (for example) by ancient population crashes, colonization dynamics, or species-specific traits and histories^8,9^. Assessment of global declines in within-population genetic diversity requires cross-generational genetic comparisons of the same population, ideally over long periods of time. Many such cross-generational studies have been performed, yet the resulting data have not been synthesised into a global estimate of the direction and magnitude of changes in genetic variation. Here we analysed temporally repeated measures of population genetic variation that were at least one organismal generation apart. We examined 5180 peer-reviewed publications and identified 76 publications on 69 species that met our criteria. These studies spanned a mean of 26 (±35 standard deviation) generations. Historical time points, defined as the age of the oldest samples, were made possible through archival specimens. These began towards the end of the industrial revolution (~1840), with the earliest sample from 1829, though the average historical sample was from 1942. Modern samples were defined as the youngest samples, and the average sample was from 2004. We compared estimates of genetic variation, namely heterozygosity and allelic diversity, across time points to assess the magnitude of genetic variation decline. We also examined factors associated with the direction and magnitude of changes observed.

We documented an average 6% (±20%) global decrease in mean expected heterozygosity (defined as the number of heterozygotes expected under Hardy-Weinberg equilibrium given the allele frequencies^10^; t=2.62, mean of differences=0.026, 95% confidence interval (CI) =0.006-0.046, df=63, p=0.011, paired t-test). A nearly identical decrease of 6% (±19%) was seen for mean allelic diversity (metrics used described in methods; t=2.59, mean of differences=0.058, CI=0.013-0.103, df=67, p=0.012, paired t-test,). In contrast to expected heterozygosity and allelic richness, observed heterozygosity did not decline significantly (t=−0.48, mean of differences=−0.006, CI=−0.029-0.017, df=54, p=0.629, paired t-test). This matches our expectations because theory predicts that, during a population decline, observed heterozygosity should be lost more slowly than expected heterozygosity^11^. Although the overall trends in expected heterozygosity and allelic diversity are consistent with previous studies, the loss of allelic diversity identified here (6%) is smaller in magnitude than the difference reported between fragmented and unfragmented natural populations (29%)^7^. The discrepancy between the two measures can be attributed to the difference in study designs: here we considered all cross-generational studies, even those where environmental conditions remained essentially constant, whereas the previous analysis examined single time-point studies and only compared populations with a known environmental difference. The decline in genetic variation identified in this study is also less severe than ongoing population declines. Wild vertebrate populations have declined by an average of 58% since 1970, at a rate of 2% per year^5^. We observed only a 6% decline in variation across an average of 62 years, thus a decline of ~2% every twenty years or 0.1% per year. This difference is anticipated because loss of genetic variation is broadly linked to genetically effective population size and not census size^10^. Importantly, historical samples are known to have a higher rate of genotyping error and smaller sample sizes, both of which can decrease estimates of heterozygosity and allelic richness^12^. Consequently a 6% decline should be viewed as conservative because historical values may underestimate original diversity.

The overall decrease in expected heterozygosity and allelic diversity suggests that rare variation has been lost —and is being lost— from contemporary populations. Because heterozygosity both prevents inbreeding depression and enables responses to selection ^e.g 1,2,13^, a 6% decline can undermine a species’ resilience to anthropogenic disturbance, including climate change. Moreover, much larger declines could be occurring in some species, geographical regions or time periods. We therefore examined what factors were correlated with the loss of genetic variation (as a proportion relative to the historic value, because relative loss is biologically important) using a generalized additive model (described in Methods; variables not discussed in the main text were not in the final model). The change in expected heterozygosity was significantly correlated with the date of the oldest sample in a study (F=15.6, df=1, p=0.0002). This relationship was non-linear, and studies using archival samples from close to the end of the industrial revolution showed a greater decline relative to those time series starting in, or after, the 1950s. After the 1950s, the observed decline in expected heterozygosity was lower, and a slight gain is possible (Fig. 1). Two species showed notable recent increases in expected heterozygosity (Fig. 1, grey circles), with these increases attributed to increased immigration^14^ and poor estimates of historical genetic diversity due to limited sample size^15^. Excluding these studies reduced the post-1950s gain in genetic diversity (Fig. S1). The number of generations between time points was also correlated with the total loss of allelic diversity, with longer studies showing a greater decline (t=2.5, df=1, p=0.019, Partial eta^2^= 0.15, linear model; Fig. S2).

**Fig. 1:**
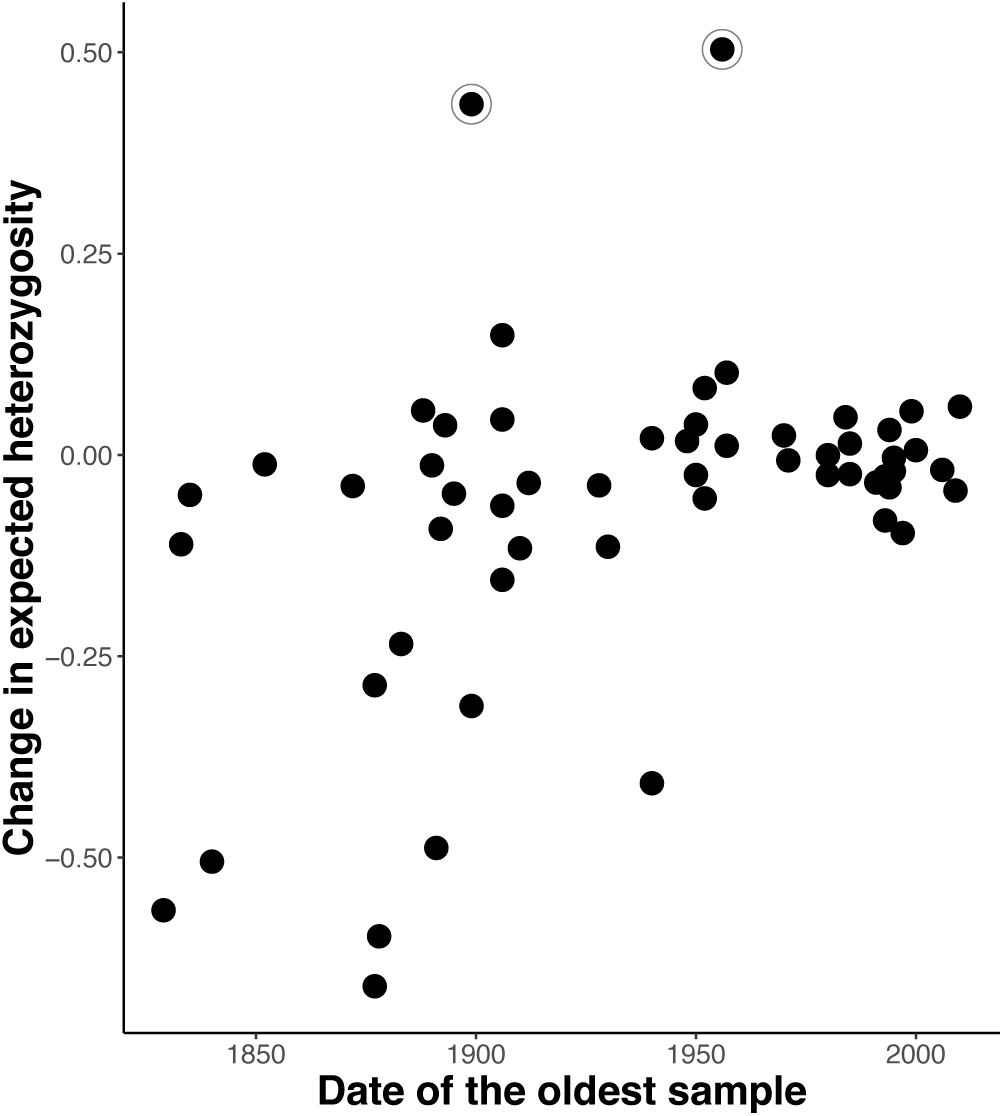
Proportional change in expected heterozygosity in modern time points (shown is the historic value subtracted by the modern, divided by the historic) relative to the age of historical samples. Circled points are two outliers where gains in variation are attributed to immigration and limited historical samples (*Capreolus capreolus* and *Atlapetes pallidiceps* ^13,14^).

Although land use change was present as early as the 1700s, fragmentation and agricultural transformation intensified dramatically after the industrial revolution^16^. Consequently, human impacts on genetic variation might have been negligible in most wild populations before the 1800s (although human mediated extinctions of megafauna would be a notable exception^17^). Estimates of variation for historical time points in the 1800s are therefore likely pre-decline and perhaps more representative of historical levels, whereas estimates are likely mid-decline for historical time points in the 1950s, when rare variation may have already been lost. These effects are likely driving the trend of reduced genetic diversity loss with younger historical samples (Fig. 1). Furthermore, the gains in expected heterozygosity after the 1950s could be due to immigration and fish stocking efforts (where populations are supplemented with captive-reared individuals), given that 13 of the 17 studies that invoke these processes have historical time points after 1950. Multi-time point comparisons of the same populations, ideally using ancient DNA to obtain estimates from before the 1700s, are needed to confirm when declines began and to estimate the total loss of genetic variation.

Post-industrial revolution human disturbances, such as land use change and habitat fragmentation, are not distributed equally across the globe and are particularly severe within Western Europe and South East Asia^16^. Furthermore population declines since the 1970s are 30% higher in neotropical regions relative to the global average^18^. We therefore hypothesized that the geographic region of a study might influence the observed loss of genetic diversity and regions of extreme change may have suffered more severe variation loss. Comparisons among continents was the finest regional scale possible for our analysis due to the limited number of studies, and continent did have a significant effect on temporal changes in expected heterozygosity (Fig. 2, F=3.7, df=6, p=0.0044, generalized additive model). Although present in the final model for temporal changes in allelic richness, pairwise comparisons between continents were all non-significant. Greater losses of expected heterozygosity were observed in Africa (z=−3.9, p=0.002, Tukey test) and North America (z=−3.2, p=0.018, Tukey test), both relative to South America (Table S1). Previous studies have found areas of above average genetic diversity within Africa^6^; however, a greater diversity loss does not appear to be due simply to a higher starting point (i.e. an initial excess of rare variation). Instead, losses in Africa are mostly driven by the severe loss of variation on islands (Mauritius and the Seychelles). Specifically, expected heterozygosity of African mainland species declined by only 0.3% on average, whereas species on African islands declined by a profound 49% on average. Considered globally, island and island-associated species generally have a greater loss of expected heterozygosity (18%, or 7% without African island species) than mainland/non-island species (2%). Island species often consist of small and isolated populations and, as a result, are more vulnerable to extreme population declines^19^. Furthermore, human impacts measured by population density, land transformation, transport infrastructure, invasive species, and electrical power infrastructure are higher on islands^19,20^. Together, these human influences are likely fuelling higher rates of fragmentation, population declines and genetic drift, which are ultimately leading to the higher loss in expected heterozygosity. Islands are hotspots of endemism^21^ and this severe genetic decline will disproportionally elevate the risk of global species loss, adding to the danger of a biodiversity extinction crisis if left unmitigated.

**Fig. 2:**
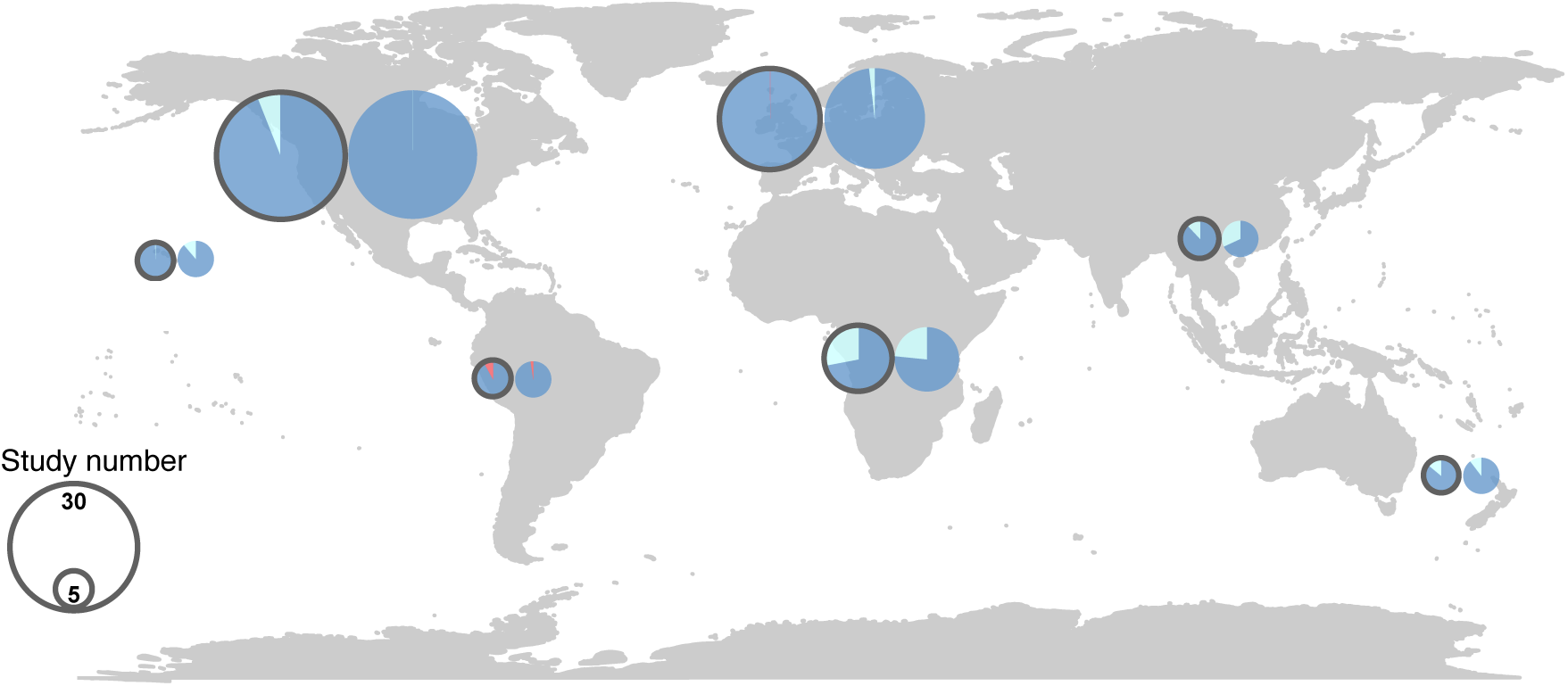
Geographic trends in genetic variation. Outlined pie charts represent expected heterozygosity and open circles represent allelic richness. Each pie chart shows the mean loss (light blue) or gain (red) of modern samples’ genetic diversity relative to the historical baseline. Pie chart size scales with the number of studies conducted on each region (regions with less than five studies are grouped into one size category).

Genotyping and sequencing efforts were globally unequal among the studies we examined, with a strong bias toward North America and Europe: 64% (n=57) of species in the dataset were from those two continents. The other continents were represented by a mean of only six studies. This regional bias is more extreme than for mitochondrial haplotype diversity estimates^2^. The geographical bias in sampling can be attributed to the rarity of historical samples, as well as the cost and specialist knowledge needed to sequence historical samples^12^. Unfortunately, the small number of studies on many continents could be biasing the trends we observed, making the conclusions we draw only provisional. South America, for example, showed an 8% gain in expected heterozygosity, but only two independent studies of five bird species were found for the region based on our search criteria. If South America is removed, continent continues to have a significant effect on expected heterozygosity in our model (F=2.9, df=5, p=0.024 generalized additive model), but pairwise comparisons show no significant differences across continents (Table S2). Immediate effort is needed to rectify the global inequality in temporal studies to better reveal the various factors influencing genetic diversity changes. To ensure cross-compatibility, future studies should ensure that the classical metrics of genetic variation (expected and observed heterozygosity, allelic diversity) continue to be reported, alongside generation times and population size estimates.

The dataset was also taxonomically biased. For instance, commercially important fish from the class Actinopterygii comprised 32% of our studies. Unsurprisingly, mammals and birds were also well-represented, encompassing 14% and 24% of studies, respectively. Worryingly, few studies on insects (n=11) or amphibians (n=3) and no studies on reptiles were found. A strong taxonomic skew towards commercially important fish, where stocking has been used over the last century, could be biasing the observed trends in genetic diversity. Thus, we also compared variation trends averaged across taxonomic classes. The previously identified trends remained reasonably stable: expected heterozygosity declined significantly by an average of 4% (Fig. 3, t=2.51, CI=0.001-0.041, mean of differences=0.021, df=6, p=0.045 paired t-test) and allelic diversity declined, though non-significantly (t=0.80, CI - 0.062-0.13, mean of differences= 0.032, df=7, p=0.449, paired t-test). As expected due to stocking activities, fish showed a smaller loss of genetic variation than other species. We recommend that the taxonomic gaps be addressed, given the importance of understanding diversity trends in, for instance, crop pollinators and non-agricultural plants.

**Fig. 3:**
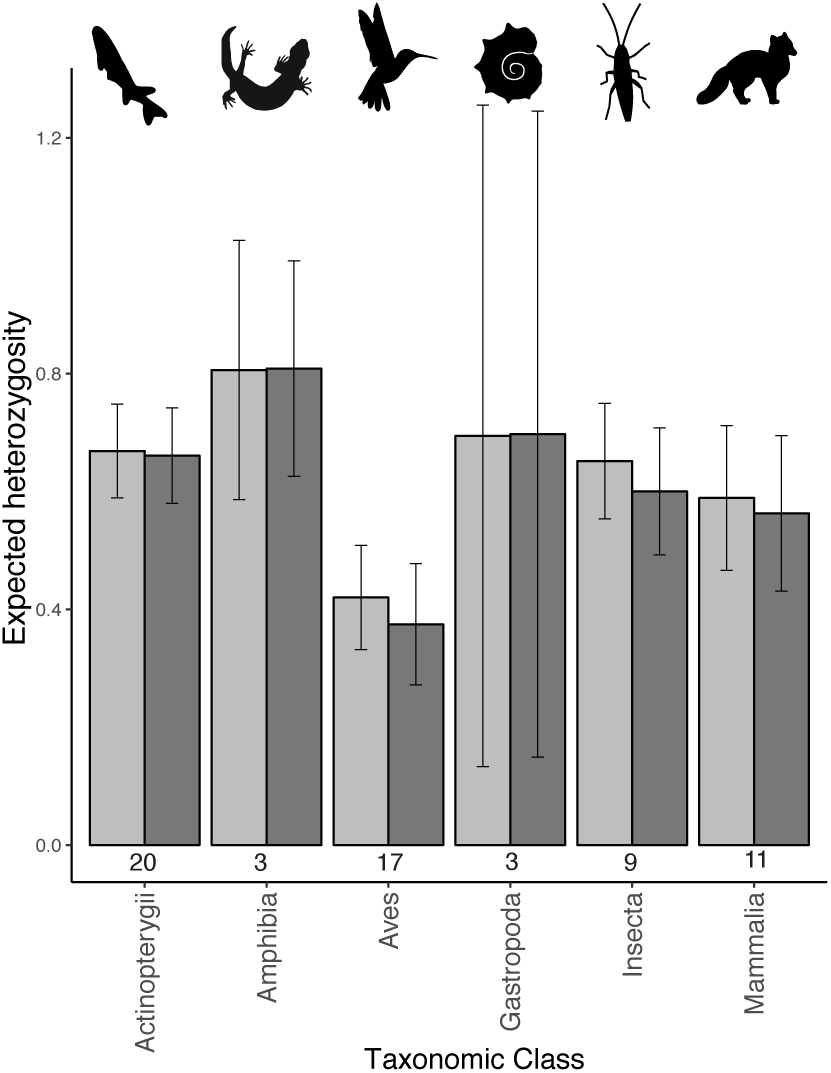
Change in expected heterozygosity across taxonomic classes as shown by means per time point (±95% confidence interval). Light grey bars represent historical samples and dark grey modern samples. Shown below each column is the number of studies for each class. Classes with only one study were excluded.

Importantly, each category of *IUCN Red List of Threated Species*^22^ was well represented across the studies (Table S3): 58% of studies were on species of least concern, meaning that the observed declines were not driven by research bias towards critically endangered species. Therefore, a decline in diversity is occurring even for species of low conservation concern. We must further stress that our requirement for modern samples (to compare diversity change through time) forced exclusion of extinct species, and thus the species that have undergone the most severe population declines. Conservative estimates suggest extinction rates are now 100 times the historical base-line rate^23^. Hence, the true total loss of genetic variation will be much greater than estimated, and a 6% decline should be viewed as conservative.

## Acknowledgments

We thank Genevieve E Finerty for her advice on the statistical analysis. We also thank Willow Rook, Erica Ponzi, Dr. Dennis Hansen and Dr. Nicolas Ponroy for their valuable comments on the text. This work was supported by a Natural Sciences and Engineering Research Council of Canada (Strategic grant to VF).

## Author contributions

DML conducted the literature and statistical analysis, and wrote the paper. AH and EVD commented on the analyses and text. VF mentored the work, and commented on the analyses and text. All authors contributed to the original idea.

## Author information

The authors declare no competing interests. Correspondence and data requests should be directed to DML.

## Methods

### Synthesis design

We searched Google Scholar (20/02/2018-24/05/2018) for peer-reviewed publications that repeatedly sampled nuclear variation in the same population at least one generation apart. Key words used were “museum sequencing”, “genetic diversity time”, “ancient DNA”, “temporal diversity genetics”, “historic specimens”, “museum specimens”, “museum specimens genomics”, “temporal genetic dynamics”, and “ghost alleles.” Metrics of genetic variation, including expected heterozygosity, observed heterozygosity and allelic diversity (as measured by allelic richness, number of alleles, nucleotide diversity [*π*], or method-of-moments inbreeding [F_H_]) were collected from 89 studies. These studies represent 82 species and 79 peer-reviewed publications, spanning an average of 55 years (±51 standard deviation) or 25 generations (±34 standard deviation). Seventy-nine species were examined with microsatellite markers, seven with single nucleotide polymorphisms, two with restriction enzyme-based markers, and one with candidate genes. Seventy-six studies spanned more than one generation. The remaining 13 studies either were too short (less than one generation) or generation time estimates were unavailable.

We estimated the mean genetic variation metrics for each of the two time points, henceforth “historic” and “modern”. Limiting the analysis to two values was necessary as many publications only had two time points. Populations in each study that were sampled in both the historic and modern time points were included in the analysis as an average per time point. Exceptions were made when population information was unavailable, in such cases values were often reported in the publications as means across all samples/populations for each time point. When populations from the historic time point went extinct before modern samples were taken, we included these extinct population’s samples in the historic time point. We recorded the length of time separating the oldest and newest sample in generations, as well as the start date of the study. If multiple time points were reported, we took the two furthest apart. If multiple estimates of generation time were given for one species, we used the median. If no generation time was given in the study reporting the genetic estimates, we used estimates for the same species from peer-reviewed literature. To account for potential research bias towards bottlenecked or inbred species, which would inflate estimates of diversity loss, we recorded whether the paper discussed if the time series began before or after a major population crash (hereafter bottleneck status), and *IUCN Red List of Threated Species* species status. When the *IUCN* ranking contradicted the study’s description, we deferred to the status in the paper. We also recorded the mean census population size in each time point, as well as the minimum number of genetic markers used. However, because population size estimates were reported in only 33 studies, this explanatory variable was excluded from our analysis. Furthermore, because genetic variation is more closely related to effective population size and not census size^10^, census size is unlikely to have an association with genetic variation. Unfortunately, effective population sizes were impossible to include due to huge confidence intervals on estimates or the use of multiple estimation methods, each giving a very different value. The country and continent of the study were recorded to test for geographic bias. Finally, to check for a bias in taxonomic representation and examine if declines were unequal across taxa, we recorded the taxonomic class of each species.

### Statistical Analysis

#### Assessing temporal trends in diversity

All analyses were conducted in R (v.3.5.0)^24^. All tests were two-sided. Paired t-tests were used to test for loss of genetic variation. Pairs consisted of the modern and historic time point values for each species in each independent study. Paired analyses were necessary because each marker set likely has different ascertainment biases and study-specific variables. Values compared were means across markers, because values for each genetic marker were often not reported. Due to multiple diversity metrics, we standardized values of allelic diversity only (allelic richness, number of alleles, *π*, or F_H_) by dividing both the historic and modern values by the historic value. This was not necessary for heterozygosity estimates because they were always reported as expected or observed heterozygosity. We tested if the mean observed or expected heterozygosity and allelic diversity differed between time points. Five species were present in multiple studies (n=13). These values were not initially excluded and were treated independently because the studies are independent and the populations were different evolutionary significant units. However, to control for potential bias due to unequal taxonomic representation, an additional test on average values per taxonomic class was conducted to control for non-independence of data.

#### Factors affecting temporal trends

We examined factors potentially influencing the change observed between time points in expected heterozygosity and allelic diversity. This was calculated by subtracting modern time point values from historic, and dividing the difference by the historic time point value. Because different allelic diversity metrics cannot be combined in this analysis, as it will lead to huge variation depending on the metric used, we constrained the analysis on allelic diversity to measures of allelic richness. The change between time points for each species was the response variable. The number of genetic markers, whether the time series began before or after a major population crash, taxonomic class of each species, continent, age of the oldest sample, and number of generations between time points were included as explanatory variables in a single model. Continent refers to the region on which or near which the focal populations are found. Due to its distance from any continental landmasses, Hawaii was considered independently. The full model contained all listed variables and no interactions because balanced replication was not present across the factors. Model selection was performed using the Akaike Information Criterion or AIC. Variables not discussed in the main text were not in the final model. An extreme outlier was removed from the analysis on allelic richness because this study described over 100 alleles at fewer than ten microsatellite loci^25^, suggesting technical errors may be present. The final model for expected heterozygosity included continent and age of the oldest sample; however, the relationship between change in expected heterozygosity and age of the oldest sample appeared non-linear. Therefore, we also fitted the model with a generalized additive model (GAM), which does not assume a linear relationship. This model had a similar AICc (GAM -AICc=27.01, linear model -AICc=27.3) and thus the values reported are from the GAM. Differences across continents were compared pairwise with a Tukey test. To correct for multiple testing, Shaffer adjusted p-values are reported.

## Data Availability

Data underlying the study, including a summary table of studies used and diversity statistics reported, will be made available on dryad upon publication. Data is available from the corresponding author should it be required for review.

---

**Figure S1:**
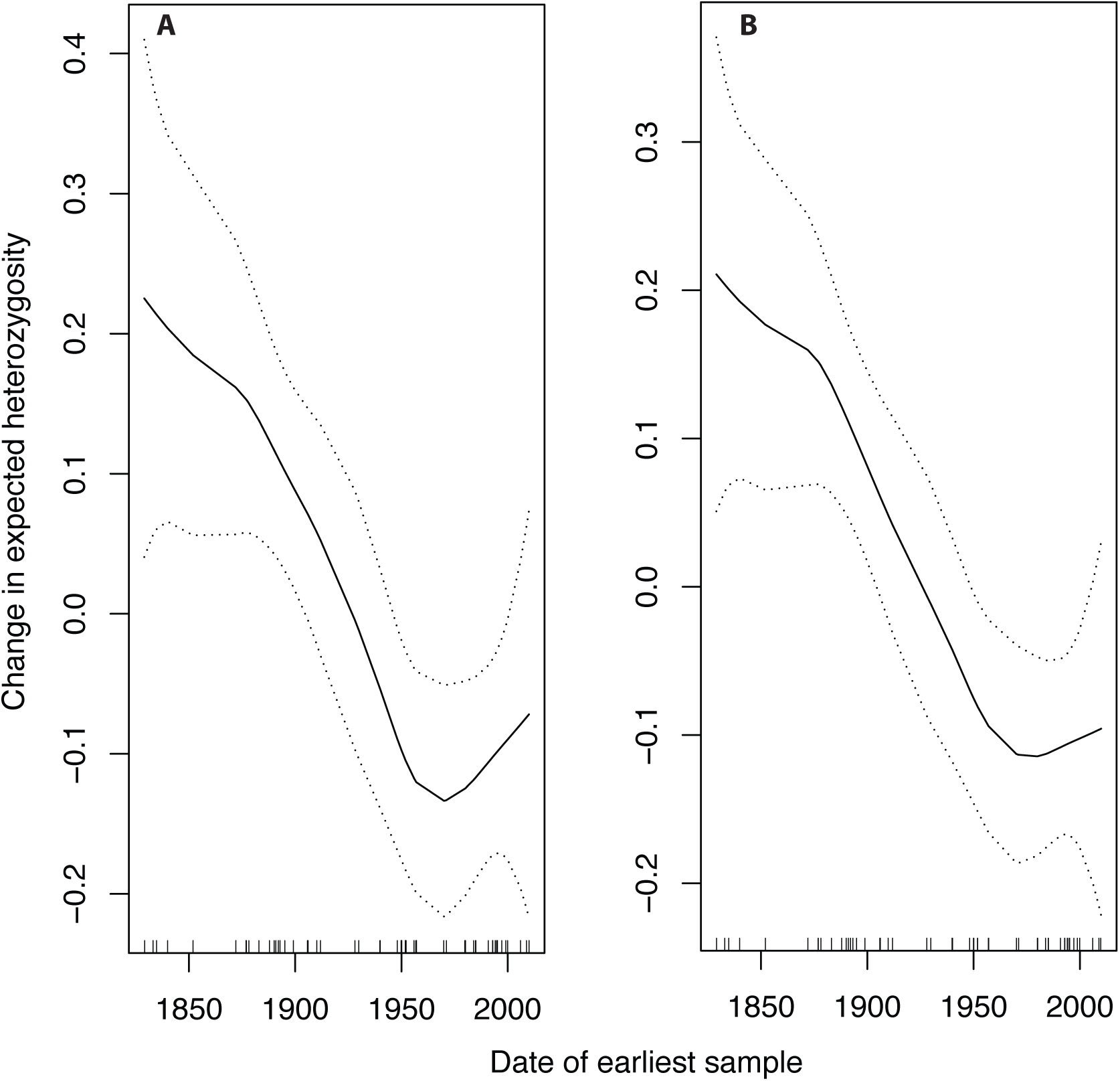
The GAM plot for the relationship between change in expected heterozygosity (a proportion relative to historic values) and the date of the earliest sample. A) Includes all data points in the synthesis and B) without two outliers that displayed high gains in expected heterozygosity due to immigration and poor historic sample size. Negative values indicate a gain in expected heterozygosity because change is estimated by subtracting modern values from historic.

**Figure S2:**
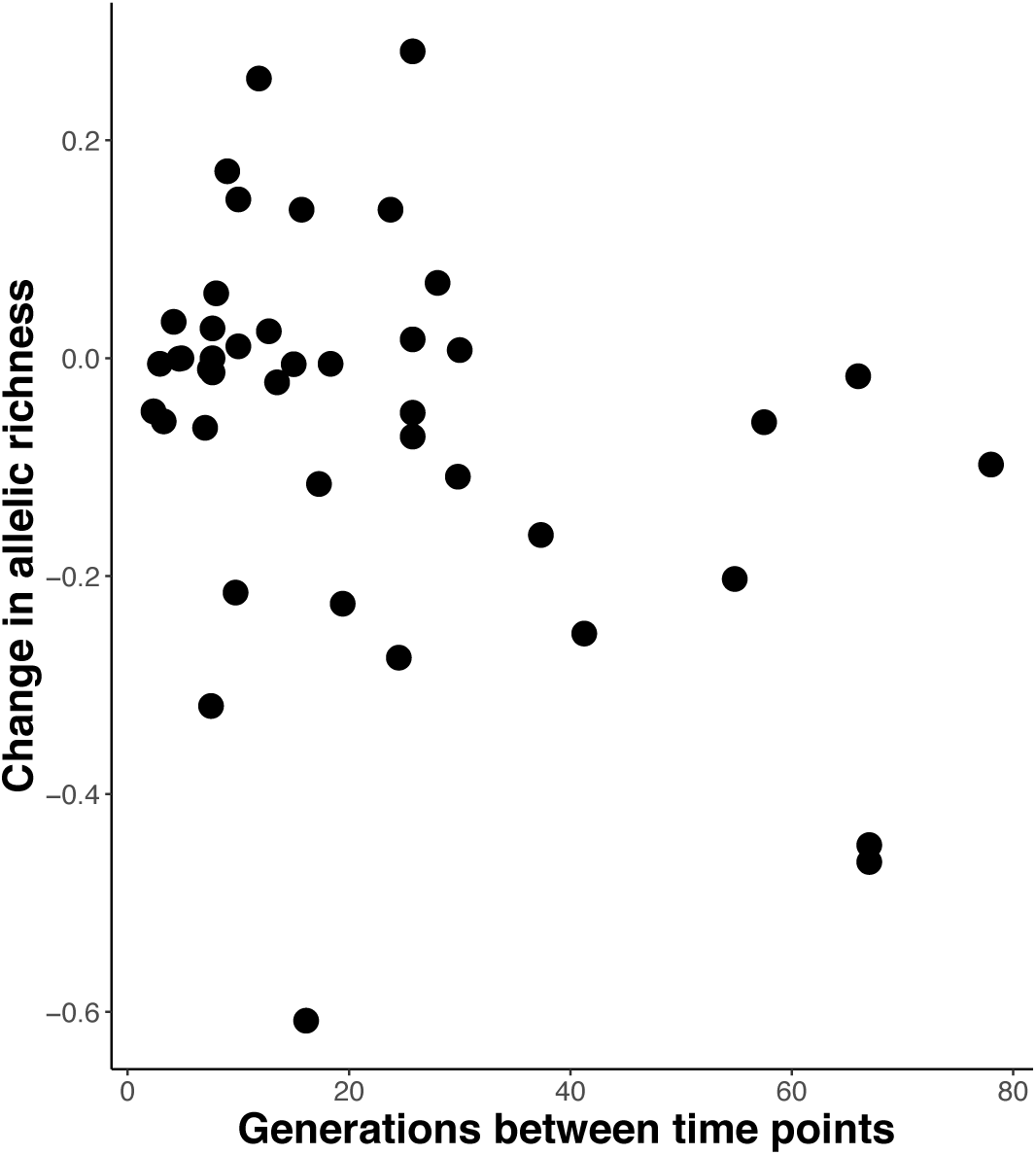
The allelic diversity (measured by allelic richness, shown is the modern value subtracted from the historic, divided by the historic value) relative to the total number of generations that a study spans.

**Table S1.** Tukey test pairwise comparisons of differences in expected heterozygosity across all continents

| Continents compared |  | Estimate of difference between means | 95% Confidence interval | Z-value | P-value |
| --- | --- | --- | --- | --- | --- |
| Asia | Africa | -0.062 | -0.398 to 0.274 | -0.54 | 1 |
| Australia | Africa | -0.175 | -0.459 to 0.108 | -1.8 | 0.503 |
| Europe | Africa | -0.198 | -0.419 to 0.023 | -2.61 | 0.101 |
| North America | Africa | -0.091 | -0.316 to 0.134 | -1.18 | 1 |
| Hawaiian islands | Africa | -0.374 | -0.767 to 0.019 | -2.77 | 0.083 |
| South America | Africa | -0.377 | -0.659 to -0.094 | -3.89 | 0.002* |
| Australia | Asia | -0.113 | -0.473 to 0.247 | -0.91 | 1 |
| Europe | Asia | -0.135 | -0.437 to 0.166 | -1.31 | 0.954 |
| North America | Asia | -0.029 | -0.331 to 0.273 | -0.28 | 1 |
| Hawaiian islands | Asia | -0.312 | -0.768 to 0.144 | -1.99 | 0.347 |
| South America | Asia | -0.315 | -0.672 to 0.043 | -2.57 | 0.114 |
| Europe | Australia | -0.023 | -0.278 to 0.233 | -0.26 | 1 |
| North America | Australia | 0.084 | -0.177 to 0.345 | 0.94 | 1 |
| Hawaiian islands | Australia | -0.199 | -0.605 to 0.207 | -1.43 | 0.916 |
| South America | Australia | -0.202 | -0.507 to 0.103 | -1.923 | 0.347 |
| North America | Europe | 0.107 | -0.059 to 0.273 | 1.87 | 0.427 |
| Hawaiian islands | Europe | -0.176 | -0.555 to 0.202 | -1.36 | 0.916 |
| South America | Europe | -0.179 | -0.431 to 0.073 | -2.07 | 0.347 |
| Hawaiian islands | North America | -0.283 | -0.668 to 0.101 | -2.15 | 0.318 |
| South America | North America | -0.286 | -0.543 to -0.029 | -3.25 | 0.018* |
| South America | Hawaiian islands | -0.003 | -0.410 to 0.405 | -0.02 | 1 |

\* indicates a p-value of <0.05

**Table S2.** Tukey test comparisons of the difference in expected heterozygosity across continents with South America excluded

| Continents compared |  | Estimate of difference between means | 95% Confidence interval | Z-value | P-value |
| --- | --- | --- | --- | --- | --- |
| Asia | Africa | -0.058 | -0.365 to 0.248 | -0.53 | 1 |
| Australia | Africa | -0.177 | -0.436 to 0.082 | -1.92 | 0.38 |
| Europe | Africa | -0.194 | -0.396 to 0.008 | -2.7 | 0.069 |
| North America | Africa | -0.086 | -0.292 to 0.119 | -1.18 | 0.948 |
| Hawaiian islands | Africa | -0.375 | -0.733 to -0.016 | -2.94 | 0.05 |
| Australia | Asia | -0.119 | -0.447 to 0.210 | -1.02 | 0.948 |
| Europe | Asia | -0.136 | -0.411 to 0.140 | -1.38 | 0.803 |
| North America | Asia | -0.028 | -0.304 to 0.248 | -0.29 | 1 |
| Hawaiian islands | Asia | -0.316 | -0.733 to 0.100 | -2.14 | 0.229 |
| Europe | Australia | -0.017 | -0.250 to 0.217 | -0.2 | 1 |
| North America | Australia | 0.091 | -0.148 to 0.329 | 1.07 | 0.948 |
| Hawaiian islands | Australia | -0.197 | -0.568 to 0.173 | -1.5 | 0.803 |
| North America | Europe | 0.107 | -0.044 to 0.259 | 1.99 | 0.274 |
| Hawaiian islands | Europe | -0.181 | -0.527 to 0.165 | -1.47 | 0.803 |
| Hawaiian islands | North America | -0.288 | -0.639 to 0.063 | -2.31 | 0.21 |

**Table S3.** IUCN Red list status of the species included in our synthesis and the total listed for all the taxonomic classes examined

| **Red list status** | **Number of temporally sampled species** | **Total species from the identified Classes of each Red List status** |
| --- | --- | --- |
| Critically Endangered | 2 | 2249 |
| Endangered | 10 | 3379 |
| Vulnerable | 11 | 4760 |
| Near Threatened | 5 | 8873 |
| Least Concern | 39 | 30254 |

